# Contribution of the Type 6 Secretion System to Apoptosis and Macrophage Polarization During *Burkholderia pseudomallei* Infection

**DOI:** 10.1101/2024.03.01.583019

**Authors:** Jacob L. Stockton, Nittaya Khakhum, Alfredo G. Torres

**Affiliations:** Department of Microbiology and Immunology, University of Texas Medical Branch Galveston, TX, 77555. USA; Department of Pathology, University of Texas Medical Branch Galveston, TX, 77555. USA

**Keywords:** *B. pseudomallei*, type 6 secretion system, macrophages, apoptosis, polarization, inflammation

## Abstract

*Burkholderia pseudomallei* (*Bpm*) is the causative agent of the disease melioidosis. As a facultative intracellular pathogen, *Bpm* has a complex lifestyle that culminates in cell-to-cell fusion and multinucleated giant cells (MNGCs) formation. The virulence factor responsible for MNGC formation is the type 6 secretion system (T6SS), a contractile nanomachine. MNGC formation is a cell-to-cell spread strategy that allows the bacteria to avoid the extracellular immune system and our previous data highlighted cell death, apoptosis, and inflammation as pathways significantly impacted by T6SS activity. Thusly, we investigated how the T6SS influences these phenotypes within the macrophage and pulmonary models of infection. Here we report that the T6SS is responsible for exacerbating apoptotic cell death during infection in both macrophages and the lungs of infected mice. We also demonstrate that although the T6SS does not influence differential macrophage polarization, the M2 polarization observed is potentially beneficial for *Bpm* pathogenesis and replication. Finally, we show that the T6SS contributes to the severity of inflammatory nodule formation in the lungs, which might be potentially connected to the amount of apoptosis that is triggered by the bacteria.

## Introduction

*Burkholderia pseudomallei* (*Bpm*) is a Gram-negative environmental saprophyte that is the causative agent of melioidosis (1). Melioidosis is a neglected tropical disease that affects an estimated 165,000 people, with approximately 89,000 deaths, a year (2, 3). *Bpm* was thought to be restricted to southeast Asia and northern Australia; however, it has been shown to have global distribution (3, 4). This geographical distribution includes the Americas where melioidosis is a recognized and growing public health threat (5–8). The clinical manifestations of melioidosis are highly variable, which often leads to misdiagnosis, earning *Bpm* the moniker “The Great Mimicker” (1, 9, 10). *Bpm* is a facultative intracellular pathogen that can successfully infect both phagocytic and non-phagocytic cell types and get distributed to almost every tissue in the host (11, 12). *Bpm* owes its success as an intracellular pathogen to an arsenal of virulence factors that it utilizes to invade, survive, and spread from cell-to-cell. One critical virulence factor is the type 6 secretion system (T6SS), *Bpm* utilizes this nanomachine to fuse host cell membranes and generate multinucleated giant cells (MNGCs) (13–16). MNGC formation is the keystone pathogenesis feature of *Bpm* and the lack of T6SS activity results in attenuation of the bacterium (13, 17). The mechanism by which host cell membranes are fused by the T6SS and the consequence of MNGC formation, from the host perspective, are both currently unknown.

Previously, we began to interrogate this intracellular pathogen by performing dual RNA-seq using our established *in vitro* model of gastrointestinal (GI) infection (18, 19). During this analysis, it was found that T6SS activity contributes to modulation of inflammatory responses through NFκB and the differential expression of numerous cell death pathway genes. The primary cell death pathway highlighted was apoptosis, which is significant due to the nature of apoptosis being “immunologically silent” (20). Cells that undergo apoptosis do not release intracellular contents that would be perceived as damage associated molecular patterns (DAMPs) by the immune system and result in an inflammatory response. Apoptotic corpses are cleared by phagocytic cells to prevent secondary necrosis in a process called efferocytosis (21). Hijacking this mechanism of cell death to evade the immune system would be advantageous to the bacterium as it can continue to spread from cell-to-cell through efferocytosis under the immune silence of apoptosis. Modulation of phagocytic cell death and evasion of intracellular clearance mechanisms by *Bpm* (22, 23) and apoptosis has been implicated during infection numerous times in a multitude of cell lines (24–28). Certain virulence factors have been implicated in the activation of apoptosis, including the type 3 secretion system (T3SS) (26) and BimA protein (24), however, the dual RNA-seq analysis implicated the T6SS apparatus (18). As the T6SS is downstream of both the T3SS and BimA-mediated actin motility during intracellular pathogenesis, it is likely that the T6SS is the main driver of apoptosis during infection.

Cell death can have a profound impact on the immune microenvironment; different modes of death can vastly change how the immune system responds. For example, pyroptosis and necroptosis release pro-inflammatory DAMPs and cytokines, which is in contrast to the silent death occurring during apoptosis (29). Both processes can impact the behavior of macrophages through the mechanism of polarization. Macrophage polarization is a phenomenon during which these immune cells get activated and skew towards pro-inflammatory (M1) or alternatively activated (M2). M2 macrophages commonly take on anti-inflammatory and homeostatic characteristics, while classically activated M1 macrophages are primarily involved in pathogen clearance and tissue damage (30). We have recently reported that *Bpm* elicits differential pulmonary macrophage polarization during infection with a *Bpm* Δ*bicA* strain, a T3SS mutant. While wild-type (WT) infected mice resulted in both M1 and M2 polarization, the Δ*bicA* failed to generate M2 polarization (31). In this work, we evaluate the contribution of the T6SS system as an inducer of apoptosis and macrophage polarization to understand the consequences of MNGC formation and start elucidating its role in pathogenesis.

## Results

### The T6SS is dispensable for survival inside of macrophages

To begin understanding how T6SS activity affects apoptosis and polarization, we first established how our T6SS mutant, Δ*hcp1* (BPSS1498), replicated inside of macrophages. Hcp1 is the most prevalent structural protein of the T6SS and hexamerizes to form the inner sheath of the injectosome, deletion of *hcp1* ablates MNGC formation while attenuating the bacterium during *in vivo* infections (13, 15). We chose to evaluate intracellular survival in two macrophage models: RAW 264.7 cells and BALB/c bone marrow-derived macrophages (BMDMs). As RAW 264.7 cells were initially derived from BALB/c mice, we chose the same background for the primary BMDMs. Unlike the previously characterized regulatory mutant Δ*bicA* (31), the Δ*hcp1* strain did not display an intracellular survival defect in either macrophage model (**Fig 1A-B**).

**Figure 1:**
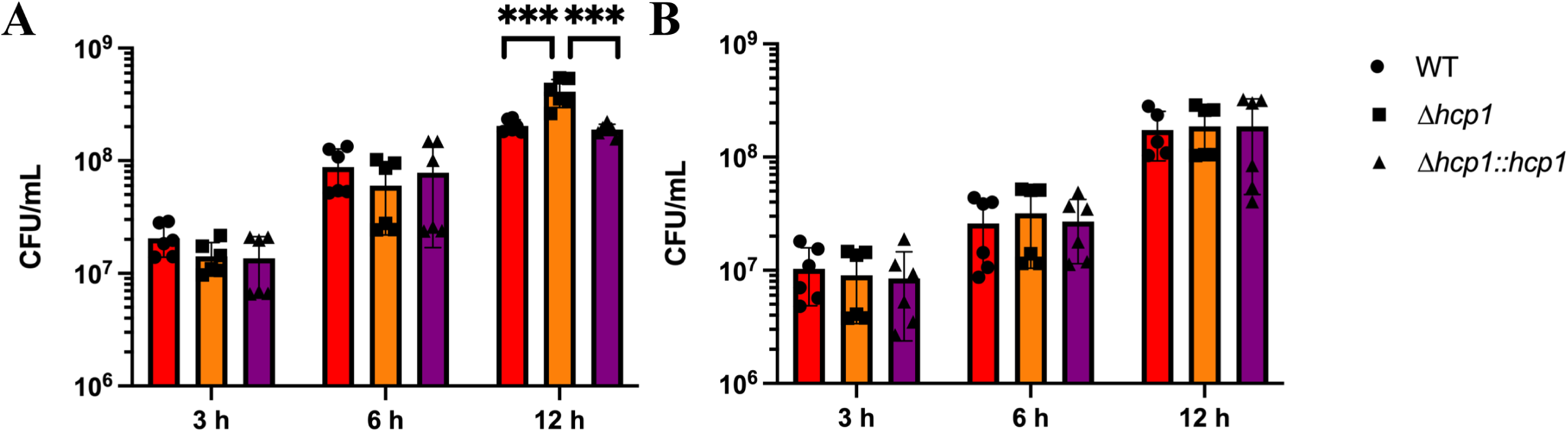
Intracellular survival of *Bpm* in macrophages is not T6SS-dependent. RAW 264.7 cells (**A**) or BALB/c BMDMs (**B**), were infected at an MOI of 10 with *Bpm* K96243 WT, Δ*hcp1,* or Δ*hcp1::hcp1* and bacteria enumerated at 3, 6, and 12 hpi to assess intracellular replication. Bars represent an average of two independent experiments performed in triplicate ± SD. Significant differences were assessed via one-way ANOVA followed by Tukey’s multiple comparison test. p < 0.05 *, p < 0.01**, p < 0.005***, p < 0.0001****.

The Δ*hcp1* strain does appear to survive significantly better than the WT or complemented Δ*hcp1::hcp1* strains in the RAW 264.7 model but the mechanism behind this phenotype remains unclear (**Fig 1A**). Together, these data suggest that although the T6SS is critical for virulence, it is dispensable for replication within macrophages.

### T6SS activity exacerbates apoptosis in macrophages and during *in vivo* infection

To evaluate apoptosis, we utilized flow cytometry to measure the externalization of phosphotidylserine (PS) via Apotracker dye paired with a live/dead (L/D) viability dye. This allows for the differentiation between apoptotic death (Apotracker +, L/D +/-) and necrotic forms of cell death (Apotracker -, L/D+). We performed an infection time course in RAW 264.7 cells and measured the percentage of macrophages that were apoptotic (Apotracker +) at 3-, 6-, 8-, and 12-hours post infection (hpi) (**Fig 2A-E**). Beginning at 6 hpi, WT infection results in significantly more apoptotic cells (**Fig 2B & E**), and by 8 hpi and 12 hpi all infection groups had high levels of apoptosis events (**Fig 2 C-E**). Although Δ*hcp1* had increased apoptosis compared to mock infected cells, WT and Δ*hcp1::hcp1* demonstrated a dramatic increase over Δ*hcp1* at both 8 and 12 hpi. High amounts of intracellular replication results in a robust apoptotic response, however, T6SS activity exacerbates apoptosis in macrophages.

**Figure 2:**
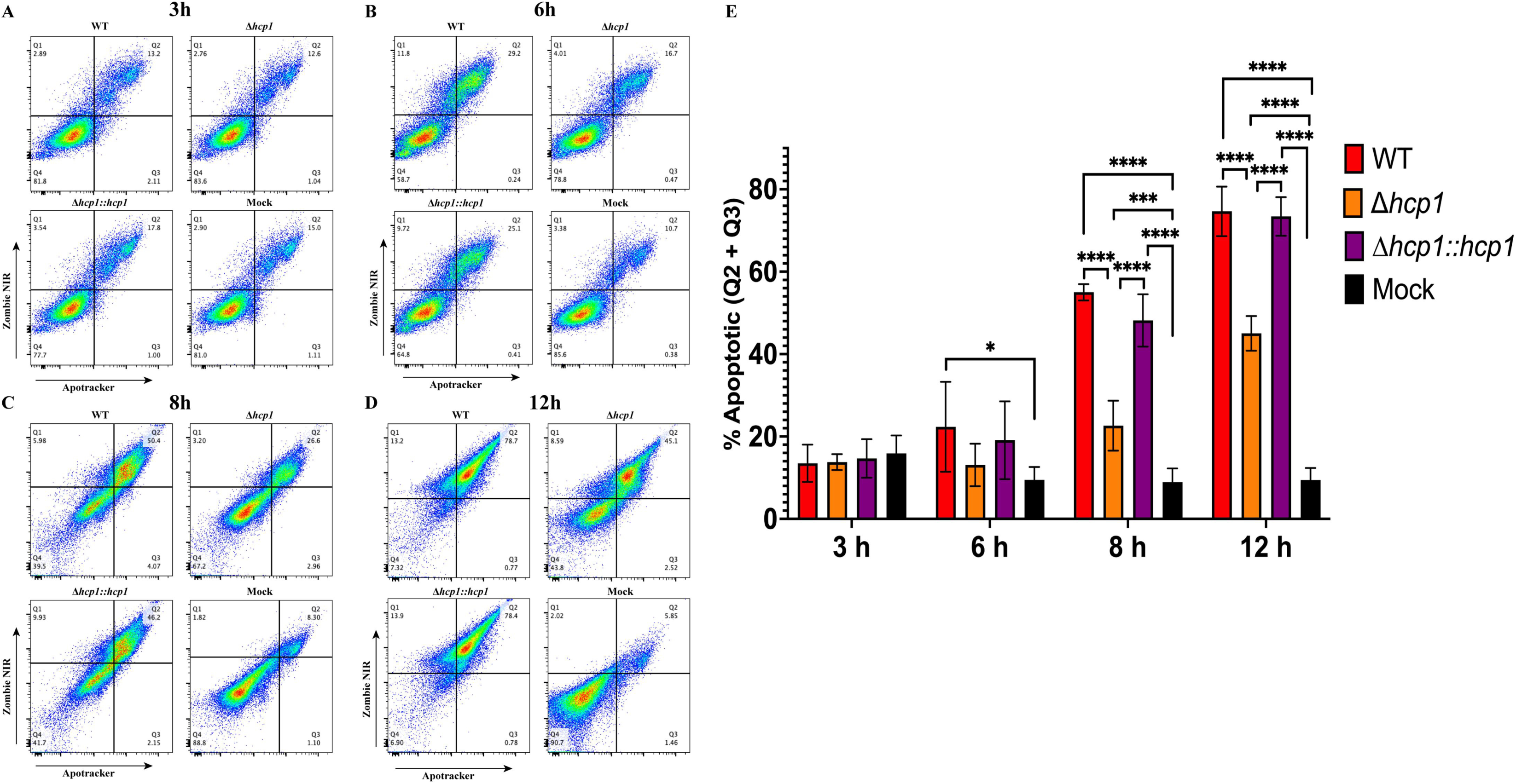
Functional T6SS exacerbates apoptosis in macrophages during infection. RAW 264.7 cells were infected at an MOI of 10 with *Bpm* K96243 WT, Δ*hcp1,* Δ*hcp1::hcp1*, or mock infected and collected at 3 (**A**), 6 (**B**), 8 (**C**), or 12 (**D**) hpi. Cells were evaluated for apoptosis via staining with Apotracker Green and Zombie NIR (Live/Dead). Percentage of apoptotic cells were counted as Apotracker+ and L/D+/- (Q2 & Q3) (**E**). Bars represent an average of three independent experiments performed in duplicate ± SD. Significant differences were assessed via one-way ANOVA followed by Tukey’s multiple comparison test. p < 0.05 *, p < 0.01**, p < 0.005***, p < 0.0001****.

With the T6SS-exacerbated apoptosis phenotype established *in vitro,* we wanted to assess apoptosis in murine lungs during pulmonary melioidosis. BALB/c mice intranasally challenged with Δ*hcp1* demonstrated 100% survival, confirming what has previously been reported (13). As expected, WT and Δ*hcp1::hcp1* challenged groups saw complete lethality (**Fig 3A**). The Δ*hcp1* infected mice exhibited minimal weight loss and were mostly clear of persistent infection on day 21 post infection (**Fig 3B & C**). We selected 48 h post infection to assess pulmonary apoptosis due to the disparity in disease severity observed between WT/Δ*hcp1::hcp1* and Δ*hcp1* at this time point. As such, another set of BALB/c mice were challenged, lungs were removed and TUNEL staining was performed to detect apoptosis. WT and Δ*hcp1::hcp1* infected lungs exhibited intense TUNEL signal, while Δ*hcp1* infected lungs displayed intermediate amounts of staining (**Fig 4**). This recapitulates what was seen *in vitro* (**Fig 2A-E**) with the Δ*hcp1* strain eliciting a small to moderate amount of apoptosis, while an active T6SS triggers large scale apoptosis.

**Figure 3:**
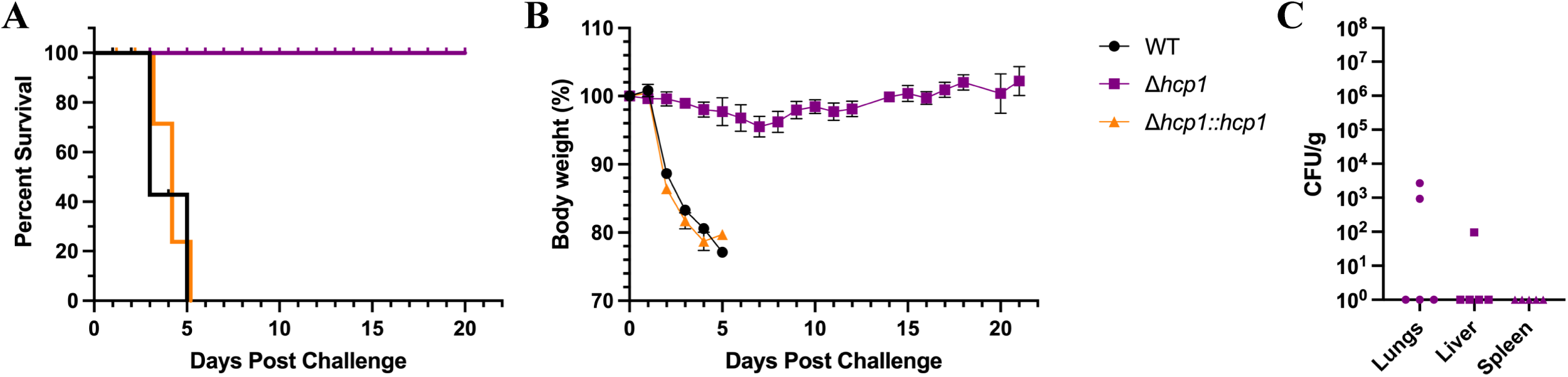
The Δ*hcp1* strain is attenuated in the intranasal melioidosis model. BALB/c mice (n = 5/group) were intranasally challenged with 3-5 LD_50_ of *Bpm* K96243 WT, Δ*hcp1,* or Δ*hcp1::hcp1* (1 LD_50_ ∼ 312 CFU) and monitored for 21 days post infection for survival (**A**) and weight loss (**B**). Animals were euthanized once the humane endpoint threshold was reached. On day 21 post infection, Δ*hcp1* survivors were euthanized and lungs, liver, and spleen were homogenized for bacterial enumeration (**C**). Error bars in (**B**) represent SEM and lines in (**C**) represent median value.

**Figure 4:**
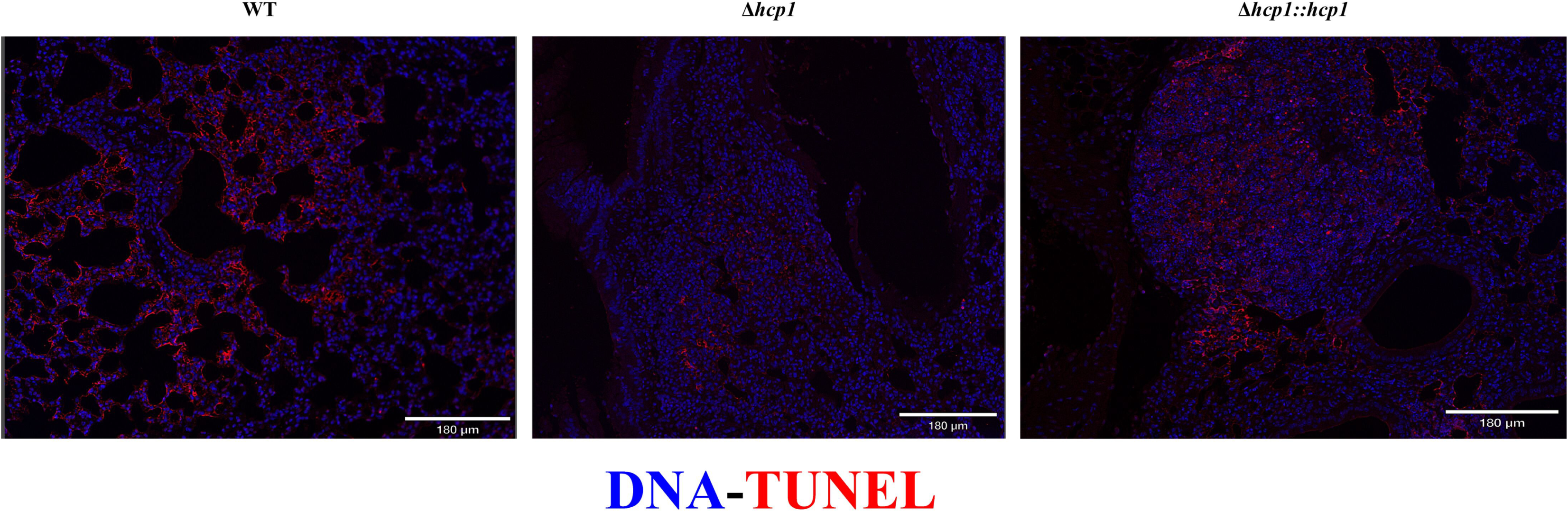
Pulmonary apoptosis mirrors *in vitro Bpm* T6SS-mediated exacerbation. BALB/c mice (n = 5/group) were intranasally challenged with 3-5 LD_50_ of *Bpm* K96243 WT, Δ*hcp1,* or Δ*hcp1::hcp1* (1 LD_50_ ∼ 312 CFU) and at 48 h post-infection lungs were harvested, formalin fixed, and mounted on slides. Sections were stained with TUNEL (red) and Hoescht 33342 (blue) to evaluate apoptosis in the lungs.

### *Bpm* infection triggers *in vitro* macrophage polarization independent of T6SS

We previously established that *Bpm* elicits both M1 and M2 polarization *in vivo* but the M2 population was BicA-dependent (31). As BicA is involved in T3SS-mediated virulence and thus the *bicA* mutant has an intracellular survival defect, we sought to understand if the differential polarization is dependent on just intracellular survival or requires a functional T6SS. Therefore, RAW 264.7 cells were infected, and at 8 hpi, were stained with markers for polarization: CD80, CD86 (M1) and CD163, Arginase-1 (M2). In this assay, cell populations that are CD80+ are being considered M1 while populations that are Arg-1+ are M2. Surprisingly, there was no significant difference in M2 polarization across the infection groups (**Fig 5A & C**) with only WT *Bpm* demonstrating an increase over mock infected cells. WT did trend higher than Δ*hcp1* on average but due to variability in the samples, this was not significant. All infection groups generated a consistent and robust M1 response (**Fig 5 B & D**). The mock infected group did exhibit a sizable amount of residual CD80 staining, however, there was a distinct shift in intensity upon infection. This result suggests that infection with an intracellular survival competent strain is enough to trigger both pro-inflammatory and alternative effector functions in macrophages and does not require of a functional T6SS.

**Figure 5:**
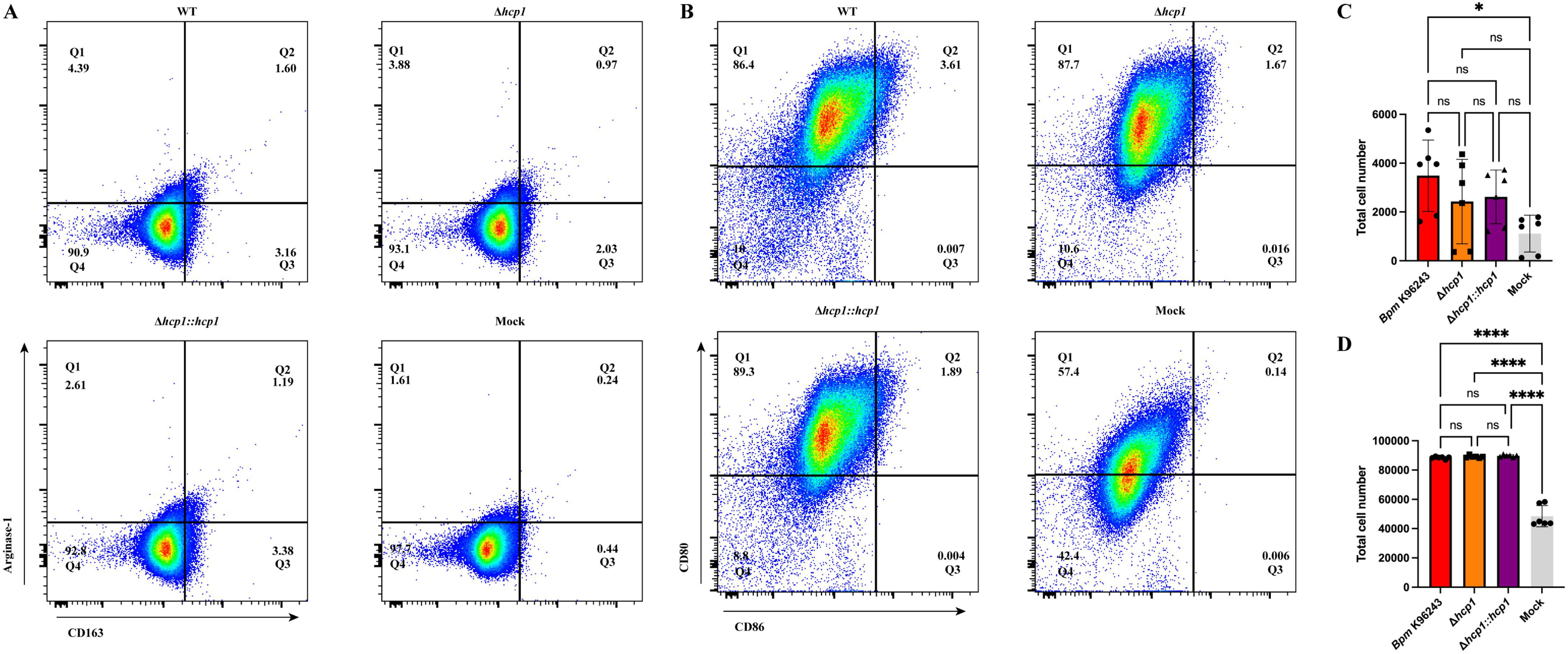
The *Bpm* T6SS does not contribute to differential macrophage polarization *in vitro.* RAW 264.7 cells were infected at an MOI of 10 with *Bpm* K96243 WT, Δ*hcp1,* Δ*hcp1::hcp1*, or mock infected and collected at 8 hpi. Cells were processed, stained, and evaluated for expression of M2 (**A & C**) and M1 (**B & D**) markers. Cells that were Arg-1+ are denoted as M2 while CD80+ cells are M1. Bars represent an average of three independent experiments performed in duplicate ± SD. Significant differences were assessed via one-way ANOVA followed by Tukey’s multiple comparison test. ns; non-significance, p < 0.05 *, p < 0.01**, p < 0.005***, p < 0.0001****.

### M2 polarization promotes intracellular survival of *Bpm*

The advantages and disadvantages of macrophage polarization for *Bpm* are unclear, as different bacterial pathogens skew polarization one way or the other to promote infection (32). To address this dichotomy, RAW 264.7 cells were pre-polarized to M1 (IFNγ + LPS) or M2 (IL-4) prior to infection and intracellular survival was assessed and compared to an M0 control (**Fig 6A-B**). M2 macrophages had decreased relative phagocytic capacity compared to M0 (**Fig 6A**) and increased intracellular survival at 3 hpi compared to both M1- and M0-polarized cells (**Fig 6B**). Interestingly, M1 polarization showed no significant advantage on bacterial clearance as compared to M0. These data further suggest that M2 skewing by *Bpm* might be offering an advantage during infection and is potentially an example of hijacking the host response to promote pathogenesis and replication.

**Figure 6:**
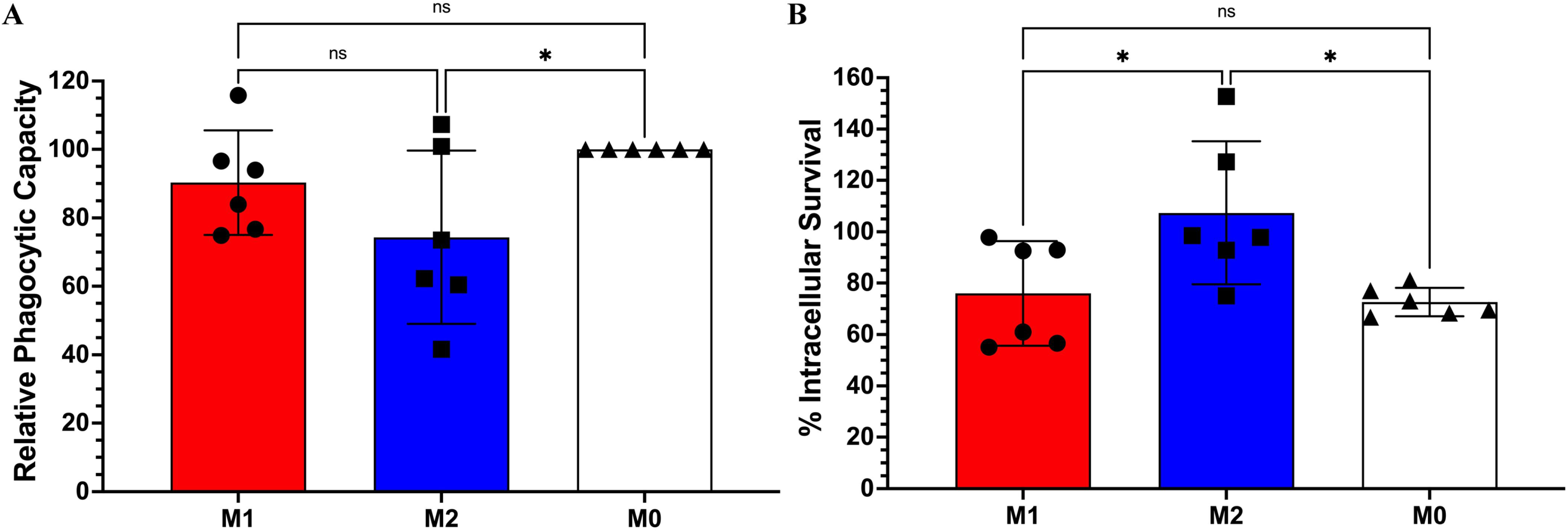
M2 polarization promotes *Bpm* intracellular survival. RAW 264.7 cells were pre-polarized with IFNγ + LPS (M1), IL-4 (M2), or media control (M0) and infected at an MOI of 10 with *Bpm* K96243. Phagocytic capacity of M1 and M2 macrophages was compared to M0 and relative phagocytic capacity was measured (**A**). Intracellular survival was evaluated at 3 hpi (**B**). Bars represent an average of two independent experiments performed in triplicate ± SD. Significant differences were assessed via one-way ANOVA followed by Tukey’s multiple comparison test. ns; non-significance, p < 0.05 ^*.^, p < 0.01**, p < 0.005***, p < 0.0001****.

### *Bpm* infection triggers *in vivo* macrophage polarization independent of T6SS

After examining the relationship between the T6SS and macrophage polarization *in vitro,* we assessed the role of the T6SS in macrophage polarization *in vivo.* BALB/c mice were intranasally challenged with WT, Δ*hcp1,* or Δ*hcp1::hcp1* and at 48 hpi lungs were removed and processed for flow cytometry. We devised a comprehensive panel (**Table 1**) and gating strategy adapted from (33) and previously utilized in (31) (**Fig S1**) to interrogate macrophage activity within the lungs. We are denoting macrophages as cells that are MHCII+ and F4/80+ after being filtered through gating and cells within that population as M1-like (CD80+ and CD86+/-) or M2-like (Arginase-1+ and CD163+). We found that although Δ*hcp1* is drastically attenuated *in vivo,* there was no difference in macrophage recruitment to the lungs during infection (**Fig 7A & B**). When examining the activation states of pulmonary macrophages, we found no difference in M1-like or M2-like macrophages (**Fig 7C & D**), however, there was a distinct downward shift in the intensity of CD80 staining in Δ*hcp1* (**Fig 7A**).

**Figure 7:**
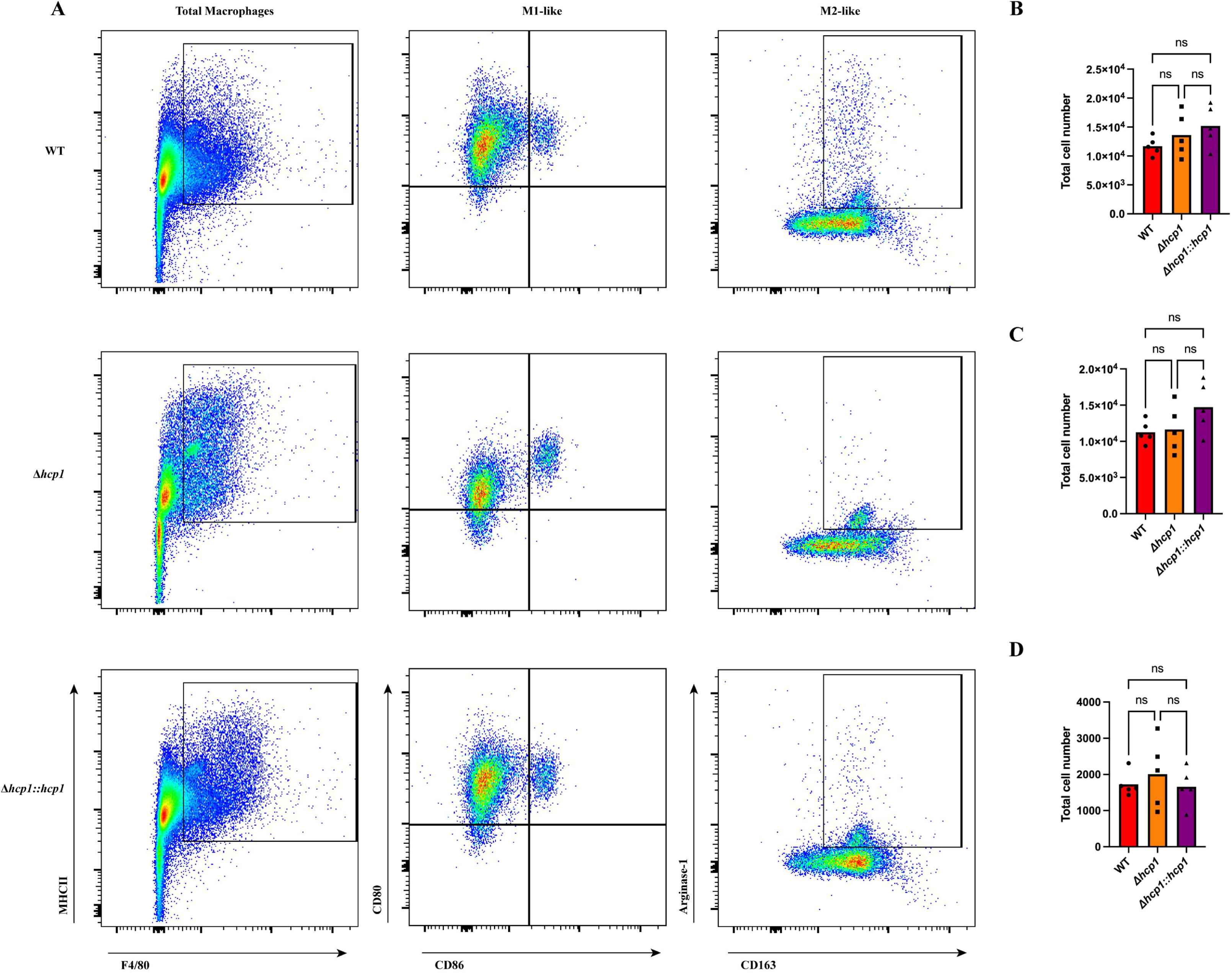
M2 polarization is not T6SS-dependent *in vivo.* BALB/c mice (n = 5/group) were intranasally challenged with 3-5 LD_50_ of *Bpm* K96243 WT, Δ*hcp1,* or Δ*hcp1::hcp1* (1 LD_50_ ∼ 312 CFU) and at 48 hpi lungs were harvested and processed for flow cytometry. A comprehensive gating strategy (**Fig S1**) was used to filter and evaluate macrophages within the lungs (**A**). Total pulmonary macrophages: MHCII+ F4/80+ (**B**), M1: CD80+ CD86+/- **(C**), and M2: Arg-1+ CD163+ (**D**) were assessed. Significant differences were assessed via one-way ANOVA followed by Tukey’s multiple comparison test. ns; non-significance. p < 0.05 *, p < 0.01**, p < 0.005***, p < 0.0001****.

**Table 1.**
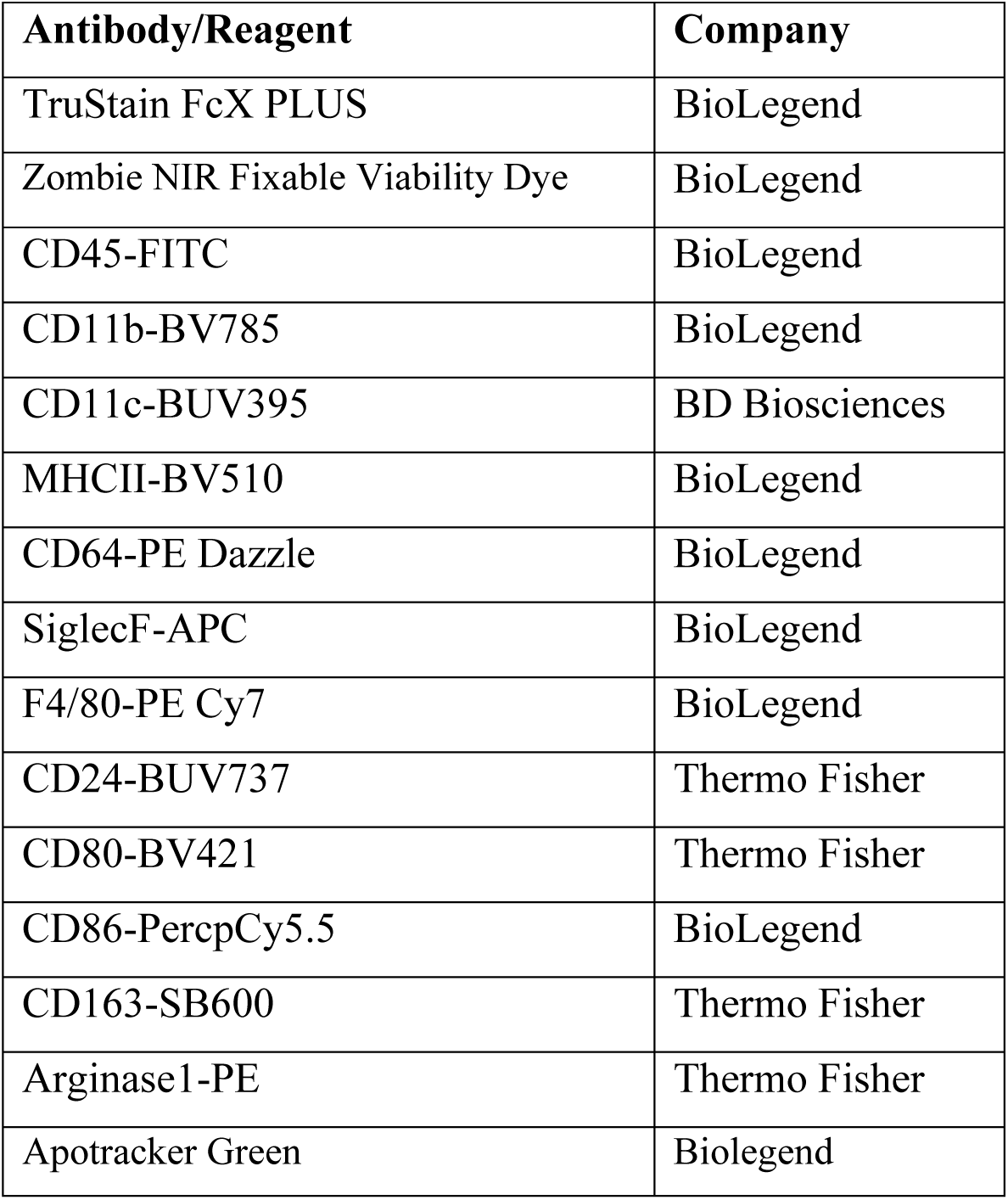
Flow cytometry antibodies and reagents.

### Inflammatory nodules predictive of M2 polarization but not T6SS-dependent

Previously, we observed that the presence of an M2 macrophage population in the lungs correlated with distinct inflammatory nodules (31). We examined H & E-stained lungs from infected BALB/c mice infected at 48 hpi for the presence or absence of this pathological feature. We found that all strains generated inflammatory nodules (**Fig 8**). However, even though the Δ*hcp1* strain generated these nodules, they were smaller and less numerous compared to WT and Δ*hcp1::hcp1* strains. The WT and Δ*hcp1::hcp1*-associated nodules appear to have more cellular debris compared to Δ*hcp1* but the cellular content of each nodule is currently unknown. We hypothesize that the nodules are likely primary replication hot spots for *Bpm* within the lungs of infected animals.

**Figure 8:**
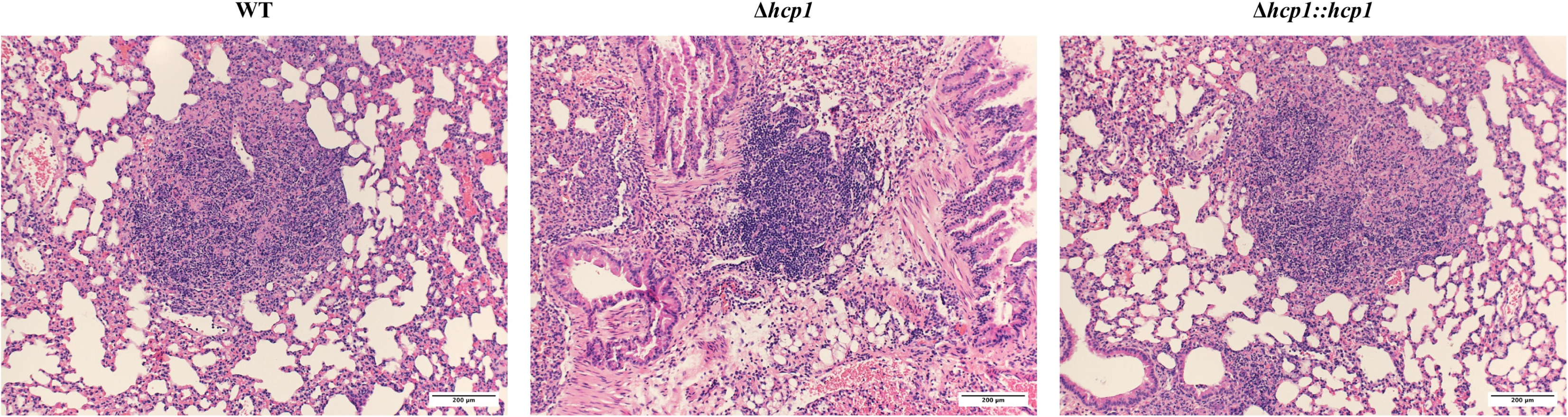
Inflammatory nodule formation is not contingent on T6SS. BALB/c mice (n = 5/group) were intranasally challenged with 3-5 LD_50_ of *Bpm* K96243 WT, Δ*hcp1,* or Δ*hcp1::hcp1* (1 LD_50_ ∼ 312 CFU) and at 48 hpi lungs were harvested, formalin fixed, and mounted on slides before hematoxylin and eosin staining. Representative images were taken using a 10x microscope objective.

## Discussion

Melioidosis is a neglected tropical disease that is a looming global public health threat (2, 3). As a facultative intracellular pathogen, *Bpm* deploys an arsenal of virulence factors to successfully survive and replicate within the host cells (1, 4). One critical virulence factor is the T6SS, an injectosome apparatus that *Bpm* utilizes to fuse host membranes and generate MNGCs. MNGC formation is the keystone pathogenesis event and T6SS mutants are highly attenuated *in vivo* (13). The mechanisms of T6SS-mediated pathogenesis and the impact that MNGC formation has on the host response is currently poorly understood. Previously, our laboratory sought to illuminate the impact of the T6SS on host response via dual RNA-seq in our *in vitro* model of GI infection (18). This analysis revealed that in the absence of the T6SS, there is substantial differential expression in pathways that are involved in inflammation, cell death, and apoptosis. The differential expression of inflammation pathways was validated by demonstrating that there is a T6SS-dependent blockage of NFκB activation, even after priming with TNFα. This work was done in primary murine intestinal epithelial cells, and it is currently unclear how this model translates to other models of infection. Therefore, to investigate how the T6SS participates in the inflammation process and cell death, we chose the *in vitro* macrophage system and the intranasal infection model to perform *in vivo* studies. Respiratory involvement is one of the most common clinical presentations of melioidosis and can progress into necrotizing pneumonia, making the intranasal model of infection particularly useful and relevant to study (1). Macrophages are a primary replicative niche for *Bpm* and are omnipresent in all host tissues, circulating or as tissue resident sentinels (11, 34). They are an attractive target for *Bpm* to manipulate as they are integral in the immune response to infection, and we have previously shown that *Bpm* is capable of differentially activating macrophages (31).

We first needed to establish how the T6SS affects intracellular survival within macrophages, as other secretion system mutants exhibit intracellular defects (18, 31, 35). We chose two different macrophage models to evaluate intracellular survival: RAW 264.7 cells, an immortalized murine macrophage cell line, and bone marrow-derived macrophages harvested from BALB/c mice. RAW 264.7 cells were initially collected from a BALB/c background, so we selected the same genetic background for our primary model. We found that, unlike the T3SS, the T6SS is dispensable for replication within both immortalized and primary macrophages (**Fig 1A & B**). There was a significant increase in intracellular survival of Δ*hcp1* in RAW 264.7 cells that was not observed in the primary BMDMs. One possibility for this result is that RAW 264.7 cells lack the inflammasome adapter protein, ASC, which limits the ability of the NLRP3 inflammasome. However, it has been demonstrated that BMDMs lacking ASC do not facilitate increased replication of *Bpm* (36).

We then evaluated the contribution of the T6SS to apoptosis of RAW 264.7 cells during infection. We found that although Δ*hcp1* triggered increased apoptosis as compared to the mock infected macrophages, WT and Δ*hcp1::hcp1* infected RAW cells exhibited very high proportions of apoptotic cells (**Fig 2A-E**). The increased viability of Δ*hcp1* at 12 hpi (**Fig 2D & E**) helps explain the increased intracellular survival in **Fig 1A** as dead and dying cells release the bacteria into the media containing kanamycin. This increase in apoptosis between 6 and 8 hpi correlates with the historical timeline of MNGC formation and thus is likely the driving force behind the rise in apoptosis. As this is an *in vitro* system, we wanted to evaluate the consistency of this phenotype *in vivo* using an intranasal challenge model using BALB/c mice. The attenuation of Δ*hcp1* has been previously documented (13), however, we needed to confirm the restoration of Δ*hcp1::hcp1 in vivo*. We found that Δ*hcp1::hcp1* recapitulates WT virulence during intranasal challenge (**Fig 3A & B**) while Δ*hcp1* remained attenuated. At day 21 post infection, the Δ*hcp1* survivors had predominantly cleared the infection, with only a couple animals harboring small numbers of bacteria (**Fig 3C**). We chose to evaluate pulmonary apoptosis at 48 hpi as there is a distinct disparity in weight loss (a predictor of disease severity) between Δ*hcp1* and WT/Δ*hcp1::hcp1* (**Fig 3B**). TUNEL staining was used to evaluate apoptosis, and paraffin embedded lung sections from 48 hpi were probed with TUNEL and a DNA counter stain and imaged to visualize relative amounts of apoptosis within the lungs (**Fig 4**). Much like the *in vitro* assay, Δ*hcp1* elicits small amounts of apoptosis but WT and Δ*hcp1::hcp1* trigger much higher and more widely distributed TUNEL signal (**Fig 4**). The *in vitro* and *in vivo* apoptosis phenotypes being nearly identical suggests that T6SS-mediated exacerbation of apoptosis is not a macrophage specific phenomenon but is a common mechanism across phagocytic and non-phagocytic cell types. There are two signaling pathways that converge on caspase-3 activation and apoptosis; the extrinsic pathway, that is initiated through an external death receptor, and the intrinsic pathway, that is triggered by internal cellular damage and release of specific mitochondrial molecules (20). The intrinsic pathway is the obvious candidate for *Bpm*-mediated apoptosis due the massive cellular trauma caused by MNGC formation. Intracellular damage, like that caused by cell fusion and ineffective ROS, is an intrinsic lethal stimulus that triggers caspase-9 mediated apoptosis. Common ligands for the extrinsic pathway are TNFα, Fas-L, and TRAIL, *Bpm* has been shown to shut down NFκB which is a driver of TNFα production. The likelihood of extrinsic activation is lower than intrinsic, but the expression of the specific molecules is unknown. However, it has also been demonstrated that splenic monocytes/macrophages from heavily colonized mice produce increased levels of TNFα and that correlated with increasing severity of pyogranulomatous lesions on the spleen (37). There is established crosstalk between the two pathways, specifically, caspase-8 cleavage of BID leading to cytochrome C release from the mitochondria and activation of the intrinsic “apoptosome”, a multimeric structure that acts as a scaffold for caspase-9 activity (38, 39). Future studies are needed to ultimately determine the signaling cascade that is involved in *Bpm*-mediated apoptosis and if there is a cell type specific contribution to the microenvironment that influences this phenotype.

We next evaluated how the T6SS affects the base inflammation state of macrophages *in vitro* via assessing the expression of polarization markers on infected RAW 264.7 cells. For this assay, an M1 macrophage is denoted as CD80+ and CD86+/- while an M2 macrophage is Arg-1+ and CD163+/-. The 8 hpi time point was selected due to the difference in the apoptosis phenotype between Δ*hcp1* and WT/Δ*hcp1::hcp1* and, although not directly measured, comparable intracellular survival. Although WT exhibited a significant increase in M2 macrophages over mock, there was no difference across the infection groups (**Fig 5A & C**). This phenotype was highly variable in the infection groups, especially within cells infected with Δ*hcp1*. On the other hand, M1 polarization was highly consistent across infection groups and all strains elicited a highly significant increase over mock (**Fig 5B & D**). Mock infected RAW 264.7 cells exhibited moderate basal CD80 expression, however, upon infection there was a distinct shift in intensity that is indicative of M1 polarization. The near complete M1 polarization tells us that the pro-inflammatory activation is the primary response to infection and that is not dependent on T6SS activity. The M2 response appears to be less evident and a secondary reaction to infection. Such activation state is highly variable and potentially is a response to the apoptosis that is occurring during infection. Traditionally, apoptotic corpses are cleared by phagocytes and anti-inflammatory molecules are released to avoid unnecessary inflammatory damage and to maintain homeostasis, a process that is called efferocytosis (21, 40). M2 polarization by *Bpm* might be incidental, an indirect response to apoptosis, but it still could be beneficial to the pathogen by creating a more permissive environment for replication. To address the question of whether macrophage polarization is beneficial to *Bpm*, we pre-polarized RAW 264.7 cells and infected with WT *Bpm* to evaluate intracellular survival. The M1 cells were pretreated with IFNγ and LPS while M2 cells were pretreated with IL-4, and expression of M1/M2 markers were validated prior to infection (data not shown). The phagocytic capacity of M1 and M2 macrophages was compared to mock treated M0 macrophages, and we found that M2s have a decreased phagocytic capacity compared to M0s (**Fig 6A**). Intracellular survival was assessed at 3 hpi and although M2s had a decreased phagocytic capacity, they facilitated increased intracellular survival compared to M1s and M0s (**Fig 6B**). This suggests that M2 polarization is a beneficial replication environment for *Bpm* and M1 polarization is inconsequential during infection. It should be noted that this phenotype is within the context of an *in vitro* infection of monocultured cells, and within the *in vivo* environment, this is more complex with multiple cell types contributing to the immune landscape during infection.

To incorporate the complexities of a multicellular immune system, we evaluated macrophage polarization in the lungs of infected mice at 48 hpi. Lungs were collected, processed, and total pulmonary macrophages and M1/M2 polarization within that population of pulmonary macrophages were evaluated (**Fig 7A**). We found that there was no difference in total macrophages present in the lungs at 48 hpi (**Fig 7B**). When we evaluated the expression of our polarization markers, we found no differences in M1 (**Fig 7C**) or M2 (**Fig 7D**) macrophages across the infection groups. This matches what was observed *in vitro* (**Fig 5A-D**). We previously reported that WT *Bpm* elicited an M2 population along with inflammatory nodules within the lungs (31) so we evaluated lung pathology at 48 hpi to determine if these inflammatory nodules were T6SS-depedendent. We found that, although they were smaller and less numerous, Δ*hcp1* infection resulted in the formation of inflammatory nodules (**Fig 8**). The structure of Δ*hcp1*-associated nodules was distinct from WT/Δ*hcp1::hcp1,* lacking the cellular debris and segmented appearance that has been observed numerous times in WT infections, both in murine and human infections (31, 37, 41). This confirms that M2 macrophages are associated with these inflammatory nodules; however, a functional T6SS might contribute to the complexity of these nodules.

Apoptotic cell death has been described as “immunologically silent” because it does not trigger a pro-inflammatory response from the phagocytic cells tasked with clearing the corpse. It has long been hypothesized that *Bpm* uses MNGC formation as a mechanism of cell-to-cell spread to prevent interacting with the external milieu and apoptosis is simply a by-product of MNGC formation. Our data begin to suggest that perhaps apoptosis is a purposeful immune evasion mechanism that *Bpm* uses to avoid triggering an effective pathogen clearing response. The robust M1 response, that occurs both *in vitro* and *in vivo,* is not effective at bacterial clearance at first glance. There is a possibility that in the absence of cell-to-cell spread and apoptosis, this M1 response is productive and can eliminate bacteria that have limited cell-to-cell mobility (Δ*hcp1)*. The correlation of M2 macrophages and the inflammatory nodule pathology might suggest that the M2 polarization is a response to tissue damage caused by the pro-inflammatory response and bacterial replication. More work needs to be done understanding which M2 subtype(s) are present inside the lung as that can shed light on their activity, as well as positioning where both M1 and M2 macrophages are in relation to the inflammatory nodules which might be working as replication hotspots.

In summary, we explored the contribution of the *Bpm* T6SS to inflammation and cell death during infection. We demonstrated that the T6SS participates as the driver of apoptosis in macrophages and within the lung but does not result in differential macrophage polarization compared to WT. The increase in inflammatory nodule severity correlated with apoptosis in the lungs, suggesting that triggering apoptosis is advantageous for pathogenesis.

## Materials and Methods

### Bacterial strains and growth conditions

All experiments were conducted with the prototypical wild-type strain of *B. pseudomallei* K96243 or derivative strains (*Δhcp1* (ΔBPSS1498) (42), *Δhcp1::hcp1*). All *Bpm* strains were routinely grown at 37°C on LB agar plates and in LB broth with shaking. *Escherichia coli* S17-1 λ*pir* were grown in LB agar plates and broth at 37°C, and Kanamycin was added for plasmid selection. For the counter selection, co-integrants were grown in YT medium supplemented with 15% sucrose.

### Construction of *hcp1* strain complementation

The *in cis* complementation of the *Bpm hcp1* mutant was performed by inserting the *hcp1* gene back into *Bpm* Δ*hcp1* strain via allelic exchange using *Burkholderia* optimized vector pMo130 (42). Purified PCR amplicon of upstream-BPSS1498-downstream and pMo130 vector were digested by NheI and HindIII restriction enzyme followed by ligation. The ligated DNA was transformed to *E. coli* S17-1 λpir donor strain. The upstream-BPSS1498-downstream/pMo130 plasmid was introduced into *Bpm* Δhcp1 strain by biparental mating as described elsewhere (18).

The clonal selection of complemented *Bpm hcp1* mutant was confirmed by PCR and sequencing at GENEWIZ.

### Macrophage culture conditions and infection assays

RAW 264.7 cells (ATCC TIB-71) were grown in Gibco Dulbecco’s Modified Eagle Medium (DMEM) plus 10% heat-inactivated fetal bovine serum (Gibco), 100 U/mL penicillin, and 100 μg/mL streptomycin (Gibco) at 37°C with 5% CO_2_. RAW 264.7 cells were maintained in T-75 flasks (Corning), detached using Accutase cell detachment solution (Biolegend) and seeded into 12 or 24 well plates (Corning). Bone marrow was collected from the femur and tibia of female BALB/c mice (Jackson Laboratories), RBCs lysed (Invitrogen 10x RBC Lysis buffer), and cells were added to polystyrene petri dishes (Sigma, 100mm x 20mm) containing RPMI 1640 w/ L-glutamine and HEPES (Gibco) plus 5 μM sodium pyruvate (Sigma), 100 U/mL penicillin, 100 μg/mL streptomycin (Gibco), 10% heat-inactivated fetal bovine serum (Gibco), and 25 ng/mL M-CSF (Biolegend). Cells were incubated at 37°C with 5% CO_2_ for 5 days with media changes on days 3 and 5. The resulting adherent cells were detached from the petri dishes using Accutase cell detachment solution (Biolegend) and seeded into 12 or 24 well plates (Corning) for further use.

RAW 264.7 cells or BMDMs were seeded at 5 × 10^5^/well in complete DMEM or RPMI without antibiotics into 24 well-plates and allowed to adhere overnight. *Bpm* strains were streaked on LB agar plates, grown at 37°C for 48 h, LB broth was inoculated and grown at 37°C with shaking for 12 h. Bacterial culture was diluted to 5 × 10^6^ CFU/mL in antibiotic free complete DMEM or RPMI and added to the cells for an MOI of 10. Cells were incubated with inoculum for 1 h for internalization, washed with PBS, and then media containing 500 μg/mL kanamycin was added to kill off extracellular bacteria. For bacterial enumeration, cells were washed twice with PBS to remove any extracellular bacteria, lysed with 0.1% TritonX-100, serially diluted in PBS, and plated on LB agar plates.

### *In vitro* evaluation of apoptosis

RAW 264.7 cells were seeded at 2 x 10^6^/well in 6 well plates and infected as described above and the infection was allowed to progress for 3, 6, 8, or 12 h. At the defined timepoint, cells were washed with PBS, removed from the well using Accustase, and pelleted in a PBS wash at 500 xg for 5 min. Cells were resuspended in 400 nM Apotracker Green (Biolegend) and incubated for 15 min before adding 1 mL of Zombie NIR (1/10,000 in PBS) for 5 min. Stained cells were washed in FACS buffer and fixed with 4% ultrapure formaldehyde for 48 h at 4°C before removal from BSL3. Analysis was done on a BD Symphony full spectrum flow cytometer. Data were analyzed using FlowJo software.

### Intranasal challenge and survival studies

Female 6–8-week-old BALB/c mice (n = 5/group) (Jackson Laboratories) were intranasally (i.n.) challenged with 3-5 LD_50_ *Bpm* K96243, Δ*hcp1,* or Δ*hcp1::hcp1* in 50 μL (25 μL/nare). One LD_50_ is equal to 312 CFU. Infected mice were monitored for survival and weight loss for 21 days post-infection and euthanized if the animal reached the threshold for humane endpoint. On day 21 post-infection, survivors were humanely euthanized, and lungs, liver, and spleen were collected for bacterial enumeration.

### TUNEL Staining

Female 6–8-week-old BALB/c mice (n = 5/group) (Jackson Laboratories) were intranasally (i.n.) challenged with 3-5 LD_50_ *Bpm* K96243, Δ*hcp1,* or Δ*hcp1::hcp1* in 50 μL (25 μL/nare). At 48 hpi, lungs were removed and fixed in 10% buffered formalin for 48 h before removal from BSL3. Lungs were sent to the UTMB Anatomical Pathology core for paraffin embedding and mounting on slides. Mounted lung sections were deparaffinized, TUNEL stained according to the included assay protocol in the Click-iT^TM^ Plus TUNEL assay Alexa Fluor 594 kit (Invitrogen), and then stained with Hoescht 33342 (ThermoFisher) to highlight nuclei. Stained slides were imaged using an Echo Revolve microscope.

### *In vitro* polarization assay

RAW 264.7 cells were seeded at 2 x 10^6^/well in 6 well plates and infected as previously described and the infection was allowed to progress for 8 h. At the end point, the cells were removed from the well via Accutase and washed in PBS before staining for flow cytometry. Briefly, cells were incubated with Zombie NIR (Biolegend) for 5 min in PBS, washed, and incubated with TruStain X plus (Biolegend) for 30 min followed by the extracellular antibodies (CD80, CD86, and CD163). Cells were fixed and permeabilized using Cytofix/Cytoperm (BD Biosciences) and stained for intracellular arginase-1. Stained cells were fixed with 4% ultrapure formaldehyde for 48 h at 4°C before removal from BSL3. Analysis was done on a BD Symphony full spectrum flow cytometer. Data were analyzed using FlowJo software.

### Pre-polarization of RAW 264.7 cells

RAW 264.7 cells were seeded at 1 x 10^6^ cells/well in 12 well plates and allowed to adhere overnight. After adherence, polarization media containing either 50 ng/mL IFNγ (MilliporeSigma) + 50 ng/mL LPS (MilliporeSigma) for M1 or 40 ng/mL IL-4 (MilliporeSigma) for M2 was added for 24 h. M0 cells were treated with mock polarization media containing an equitable amount of DMSO for 24 h. This protocol was validated via flow cytometry (data not shown) before progressing to infection assays.

### Flow cytometry

Female 6–8-week-old BALB/c mice (n = 5/group) (Jackson Laboratories) were i.n. challenged with 3-5 LD_50_ *Bpm* K96243, Δ*hcp1,* or Δ*hcp1::hcp1*; at 48 hpi, animals were euthanized, and lungs harvested for processing. Lung tissue was cut into small pieces and dissociated via incubation for 30 min at 37°C with slight rocking in RPMI plus 0.5 mg/mL collagenase IV and 30 μg/mL DNase I. The dissociated tissue was homogenized through a 100 μm cell strainer and fibroblasts and debris was pelleted via a 60 xg centrifugation for 1 min. Supernatant was collected and RBCs were lysed for 5 min at RT. Following washes, pulmonary cells were adjusted to 1 x 10^6^ cells and stained using the reagents in Table 1. Briefly, cells were incubated with Zombie NIR (Biolegend) for 5 min in PBS, washed, and incubated with TruStain X plus (Biolegend) for 30 min followed by the extracellular antibodies (Table 1). Cells were fixed and permeabilized using Cytofix/Cytoperm (BD Biosciences) and stained for intracellular markers. Fully stained cells were resuspended in 4% ultrapure formaldehyde in PBS for 48 h in accordance with the inactivation protocol approved by UTMB Department of Biosafety before removal from BSL3 laboratory for analysis via BD Symphony full spectrum flow cytometer. Data were analyzed using FlowJo software.

### Lung pathology

Lungs were collected from mice after humane euthanasia 48 h post-infection and fixed in 10% formalin for 48 h. Formalin fixed lung samples were submitted to the UTMB Anatomical Pathology core for paraffin embedding, mounting, and H&E staining. Slides were imaged using an Olympus BX51 microscope.

### Ethics statement

All manipulations of *B. pseudomallei* were conducted in CDC/USDA-approved and registered BSL3 facilities at the University of Texas Medical Branch (UTMB) in accordance with approved BSL3 standard operating practices. The animal studies at UTMB were carried out humanely in strict accordance with the recommendations in the Guide for the Care and Use of Laboratory Animals by the National Institutes of Health. The protocol (IACUC no. 0503014E) was approved by the Animal Care and Use Committee of UTMB.

### Statistical analysis

All statistical analysis was done using GraphPad Prism software (v9.0). P-values of < 0.05 are considered statistically significant. Survival differences were assessed via Kaplan-Meier survival curve followed by a log-rank test. An ordinary one-way ANOVA followed by Tukey’s post hoc test was used to analyze differences in intracellular replication and flow cytometry populations.

## Acknowledgments

This work was funded by USDA APHIS AP20VSD&B000C087. JLS is supported by a USDA APHIS NBAF Scientist Training Program Fellowship. We would like to thank Meredith Weglarz in the UTMB Flow Cytometry Core for the expertise and help in designing and implementing the flow cytometry experiments. We would also like to thank Alex Badten for his help during animal experiments, Dr. Alison Coady for allowing us to utilize her Echo Revolve microscope, and Paige Diaz for training and troubleshooting on the Echo microscope.

